# SPE-CZE-MS quantifies zeptomole concentrations of phosphorylated peptides

**DOI:** 10.1101/2024.12.07.627347

**Authors:** Lia R. Serrano, J. Scott Mellors, J. Will Thompson, Noah M. Lancaster, Margaret Lea Robinson, Katherine A. Overmyer, Scott T. Quarmby, Joshua J. Coon

**Author notes:** To whom correspondence should be addressed: Department of Biomolecular Chemistry, 440 Henry Mall, Room 4416, Madison, WI 53706.

## Abstract

Capillary zone electrophoresis (CZE) is gaining attention in the field of single-cell proteomics for its ultra-low-flow and high-resolution separation abilities. Even more sample-limited yet rich in biological information are phosphoproteomics experiments, as the phosphoproteome composes only a fraction of the whole cellular proteome. Rapid analysis, high sensitivity, and maximization of sample utilization are paramount for single-cell analysis. Some challenges of coupling CZE analysis with mass spectrometry analysis (MS) of complex mixtures include 1. sensitivity due to volume loading limitations of CZE and 2. incompatibility of MS duty cycles with electrophoretic timescales. Here, we address these two challenges as applied to single-cell equivalent phosphoproteomics experiments by interfacing a microchip-based CZE device integrated with a solid-phase-extraction (SPE) bed with the Orbitrap Astral mass spectrometer. Using 225 phosphorylated peptide standards and phosphorylated peptide-enriched mouse brain tissue, we investigate microchip-based SPE-CZE functionality, quantitative performance, and complementarity to nano-LC-MS (nLC-MS) analysis. We highlight unique SPE-CZE separation mechanisms that can empower fit-for-purpose applications in single-cell-equivalent phosphoproteomics.

## INTRODUCTION

Mass spectrometry (MS)-based proteomics is greatly benefited by the common use of chromatographic separations such as liquid chromatography (LC). In addition to chemical separation, chromatography concentrates analytes sharing a particular biochemical property, such that the chances of detecting and reliably measuring the analyte are maximized. Another separation technology that has been used in tandem with mass spectrometry is capillary zone electrophoresis (CZE). Aspects of CZE position it well for highly sensitive mass spectrometric analysis of trace sample amounts. First, the analytical volumes typically employed are relatively low (tens of nanoliters or less). The resultant flow rate is about half that of typical nano-LCMS (nLC-MS) regimes, and perhaps benefitting from the resultant sensitivity gains from improved desolvation.^1,2^ Further, column volumes for CZE are typically about five times less than that for nLC, thus creating less opportunities for sample adsorption. Finally, typical peak widths generated by CZE are relatively narrow. Given a high enough concentration, the analyte signal concentrated within such a tight temporal band should theoretically generate higher-intensity peaks than would bands with a greater temporal spread.

One limitation of CZE is the loading volume restriction that hinders achievable MS detection sensitivity, especially for complex mixtures for which the electrospray ionization (ESI) ion current is divided by a mixture of analytes at any given time. The loading amount restriction associated with CZE devices is largely due to the low-volume separation channel. If too much sample volume is loaded onto the channel, the remaining channel distance may be too short to perform an effective separation.^3^ Small diameters are desirable to ensure maximal radial uniformity of the analytes such that the migration velocity remains consistent for a given analyte. Migration velocity can be affected by a given analyte’s radial position within the separation channel because of the temperature gradient that exists, with temperature proportional to the distance from the channel walls. Since heat dissipates through the channel walls, larger channel volumes are more vulnerable to these effects. Not only is the retarded heat dissipation in radially-larger channels exacerbating effects from Joule heating, but so too is the larger amount of heat that is generated relative to lower-volume channels.^4,5^ Further, fluid and mass transfer of sub-microliter volumes is challenging and an active area of innovation.^6^ Because of these sample volume limitations, strategies to concentrate the sample pre-separation are common practice.^7–14^

Chip-based CZE devices offer a streamlined format for precise and rapid liquid handling of low volumes^15^ and are highly amenable to commercialization. Coupling these devices with electrospray ionization for mass spectrometric analysis has posed various challenges. Proper geometries are required to facilitate aerosolization of analyte ions, which drove many to integrate nanospray emitters onto the microfluidic chip. This strategy contrived additional junctions, creating vulnerabilities associated with non-closed systems such as dead volumes preventing maximum sample transfer. In 2008, Mellors *et al*.^16^ described a chip-based CZE device equipped with a fully-integrated electrospray ionization emitter, enabled by an innovative chip geometry to facilitate electrospray. This device has been commercialized (ZipChip, 908 Devices) and used for various applications.^17–24^

More recently, this device has been modified to integrate a 1 mm solid-phase extraction (SPE) bed composed of 5 μm C18 particles for pre-concentration before electrokinetic focusing and separation.^8,25^. An advantage of this technique over other common pre-concentration strategies is its higher sample volume loading capacity. This benefit is inherent to the mechanism of capturing the analyte on the SPE bed, as many channel volumes can flow through the device, whereas electrokinetic based focusing strategies are still ultimately limited by the dimensions of the separation channel. Thus, the only limiting factors for SPE-based concentration is the dissociation rate of the analyte from the solid phase material impacted by loading volume and the loading capacity of the stationary phase.

For chromatography-based separations, although the peaks generally become narrower as you increase flow rate, and perhaps temporally condensing the ion packet such that its MS signal is more intense, the achievable separation resolution (theoretical plate distance) decreases linearly.^26^ This differs from CZE because as you increase the electric field, thus expediting the separation, the widths of the peaks compress to a greater extent than does the entire separation window, therefore effectively increasing peak capacity (number of theoretical plates is not proportional to length of the separation channel or time of analysis) (see **Eqn. 1**).^27^ Therefore, achieving a narrower analyte peak using CZE could grant the added advantage of increased sensitivity while also enhancing the peak capacity.

To take advantage of the fast and consequently high-resolution electrokinetic separations CZE can offer, compatibility with MS is required to effectively acquire quantitative -omics data. Complex mixtures separated in a relatively short period of time (<30 min) challenge the sensitivity of the MS detector and the mass resolution of the mass analyzer. Further, extracted ion-based quantification is contingent upon sufficient MS sampling rates to allow reliable peak area integration.^28^ The recently released Orbitrap-Astral mass spectrometer (Thermo Fisher Scientific) possesses crucial characteristics fit for the task of maximizing the gains from SPE-CZE separation paired with MS analysis. Namely, fast sampling rates are achieved through a high degree of parallelization and a sensitive detector enabling high resolution accurate mass (HRAM) analysis. The architecture of the OT-Astral enables ion accumulation before mass analysis in the Astral analyzer, improving signal to noise ratios which are otherwise achievable through temporally expensive spectral averaging.^29–36^

CZE-MS has been demonstrated to effectively analyze single-cell proteomics samples.^37–41^ Analysis of phosphoproteomics from single to tens of cells is even more challenging when considering that phosphorylated peptides represent a fraction of the total starting material and require additional sample preparation steps that can result in further sample loss. The gold standard technology used for phosphoproteomics analysis is nano-flow LC-MS/MS. CZE-MS has been demonstrated to facilitate the identification of a complementary population of phosphorylated peptides, but primarily within bulk material or across tens of minutes of separation time.^11,42–45^ Here, we explore fast (< 15 min) separations for single-cell phosphoproteome-equivalent sample amounts (i.e. ∼1-5% of typical single-cell protein recovery) on an SPE-integrated CZE device (908 Devices, Boston MA) coupled to the Orbitrap Astral mass spectrometer. Using a set of 225 phosphorylated peptide standards we define specific chemical biases that enable CZE detection of a population of phosphorylated peptides distinct from nLC MS-derived identifications. Further, we extend our analysis to phosphorylated peptide-enriched mouse tissue to characterize the performance on complex mixtures. We report reliable quantification down to 640 zmol amounts for ∼60% of our synthetic phosphorylated peptide standards. Further, we highlight a SPE-CZE-specific advantage on the metrics of selectivity, detectability, and sensitivity for a distinct set of phosphorylated peptides.

## METHODS

### Mouse tissue lysis and digestion

Mouse brains were pulverized under liquid nitrogen, and then weighed out on dry ice for lysis. For lysis, 5.4 M guanidine hydrochloride (Sigma, 8 M, pH 8.5, G7294-100ML) in 100 mM Tris buffer (Invitrogen, 1 M Tris pH 8.0, 0.2 µm filtered, AM9856) was added to the brain tissue samples to give ∼15 mg tissue/mL. The samples were sonicated in a bath sonicator chilled to 8°C for 10 min. A methanol precipitation was performed by adding methanol (Fisher Scientific, Optima LC/MS) to 90% (v/v), vortexing to mix, and centrifuging at 9000 g for 5 min at 4°C. The supernatant was discarded and the protein pellet was allowed to air dry for 5 min prior to resuspending in 8 M Urea (Sigma, BioReagent, for molecular biology, U5378) in 100 mM Tris buffer (Invitrogen, 1 M Tris pH 8.0, 0.2 µm filtered, AM9856) with 10 mM TCEP (Sigma-Aldrich, C4706-2G) and 40 mM chloroacetamide (Sigma-Aldrich, ≥ 98%, C0267-100G) at a target concentration of ∼1 mg/mL protein (based on an estimate of 1 mg protein per 15 mg brain tissue). LysC (VWR, 100369-826) was added at a 1:50 enzyme:protein ratio and the samples were gently rocked for 4 hr at ambient conditions, followed by dilution to 2 M urea with 100 mM Tris, pH 8, trypsin (Promega, Sequencing Grade Modified Trypsin, V5113) addition at 1:50 enzyme:protein ratio and overnight incubation. To stop the digestion, 10% TFA (Sigma-Aldrich, HPLC grade, >99.9%) was added to lower the pH below 2. The samples were centrifuged at 9000 g for 5 min and then the samples were desalted using Strata-X 33 μm polymeric reversed phase SPE cartridges. Desalted samples were dried down in a SpeedVac (Thermo Scientific) and stored at -80°C prior to phosphorylated peptide enrichment.

### Phosphorylated peptide enrichment

Samples were resuspended in 0.2% formic acid (Fisher Scientific, LC-MS grade) in water (Fisher Scientific, Optima LC/MS) and the peptide concentration was measured with a NanoDrop spectrophotometer (Thermo Scientific). The samples were then dried down and resuspended in 80% acetonitrile (ACN) (Fisher Scientific, Optima LC/MS)/6% trifluoroacetic acid (TFA) (Sigma-Aldrich, HPLC, >99.0%) and vortexed thoroughly to resuspend. A 100 μL volume per enrichment tube of MagReSyn Ti-IMAC beads (ReSyn Biosciences, MR-TIM010) were washed three times with 80% ACN/6% TFA. The samples were loaded onto the beads (∼1 mg peptide per tube) and vortexed for 60 min at ambient conditions. The supernatant was removed and the beads were washed 3 times with 1 mL 80% ACN/6% TFA, 1 time with 1 mL 80% ACN, 1 time with 1 mL 80% ACN/ 0.5 M glycolic acid (Sigma-Aldrich, 99%, 124737-500 G), 3 times with 1 mL 80% ACN, and finally eluted with 2 x 300 μL washes with 50% ACN/1% ammonium hydroxide (Sigma-Aldrich, 28% in H2O, ≥99.99% trace metals basis). The samples were then acidified with the addition of 15 μL of 10% TFA. Samples were dried down with a SpeedVac (Thermo Scientific) and desalted as described previously. The dried desalted samples were resuspended in 0.2% formic acid and the phosphorylated peptide concentration was estimated using a NanoDrop spectrophotometer.

### Phosphorylated peptide standard dilution

Phosphorylated peptide standards from SpikeMix PTMKit 52 (JPT, SPT-PTM-POOL-Yphospho-1), SpikeMix PTM-Kit 54 (JPT, SPT-PTM-POOL-STphospho-1), MS PhosphoMix 1 (Sigma-Aldrich, MSP1L-1VL), MS PhosphoMix 2 (Sigma-Aldrich, MSP2L-1VL), and MS PhosphoMix 3 (Sigma-Aldrich, MSP3L-1VL) were resuspended in 0.2% formic acid, 20% acetonitrile, 80% water and pooled into an equimolar mixture of 225 phosphorylated peptide standards (**Supplementary Data Table 1**). The mixture was then diluted in sample diluent (0.01% nonaethylene glycol monododecyl ether, 100 mM ammonium acetate, 1% acetonitrile) to 1 fmol/μL then serially diluted five-fold down to 0.512 zmol/μL. For nLC-MS experiments the same equimolar mixture was serially diluted in 0.2% formic acid to match the concentration curve described for SPE-CZE analysis.

### nLC-MS

2 μL of sample was loaded onto a nano-capillary column (75 µm i.d., 363 µm o.d., Molex) etched and packed in-house at 30,000 psi^46^ with 130 A pore size Bridged Ethylene Hybrid C18 (Waters Corporation) to 30 cm. Peptides were chromatographically separated for 20.1 min using a reversed-phase gradient from 6-60% mobile phase B (MPB, 80% acetonitrile/0.2% formic acid) flowing at 300 nL/min and held at 55°C. From 0-2 min, MPB increased 0 to 6%. Then from 2-16 min, MPB increased from 6 to 60%. MPB was then held at 100% for 4 min before column re-equilibration for 16 min at 100% mobile phase A (MPA, 0.2% formic acid). Peptides eluting from the column were electrosprayed into an Orbitrap Astral mass spectrometer at 2 kV with respect to ground. Precursor ions ranging from 380-980 m/z were measured in the Orbitrap analyzer at a resolving power of 240,000, requiring either 3,000,000 charges or a 10 ms maximum injection time for each scan. Precursors from 380-980 m/z were selected for MS2 using data-independent analysis (DIA) and 2 Th quadrupole isolation windows (299 total windows across the mass range). Normalized collision energy (NCE) was set to 27%. Fragment ions from m/z 150-2,000 were measured in the Astral analyzer, requiring either 50,000 charges or a 3.5 ms maximum injection time.

### SPE-CZE-MS

For each analysis, the SPE bed was conditioned with weak mobile phase (100 mM ammonium acetate, 1% acetonitrile) before sample injection. Sample was then loaded onto the SPE bed for 240 seconds (∼ 2 μL total volume). Peptides were eluted from the SPE bed with background electrolyte (BGE, 50% acetonitrile, 1% formic acid) and shuttled to the 22 cm separation channel (70 µm x 10 µm, ∼ 30 µm i.d.) using pressures applied to the sample well and an upstream BGE reservoir. After electrokinetic focusing via transient isotachophoresis, capillary zone electrophoresis was performed using a 500 V/cm electric field between 15 kV (anode) and 2.4 kV (cathode) electrodes (**Figure 1A, Supplementary Figure S2**) for 15 min. Pressure is applied to the sample well and the BGE well during elution with the strong mobile phase and again after 6 min of separation to assist the lower-mobility analytes. Pressure is also applied to the cathode (HV2, **Supplementary Figure S2**) to assist electrospray ionization. Gas is supplied by the Aux and Sheath gas lines from the Orbitrap Astral. These pressures were set to maximize sample transfer from the SPE bed to the separation channel and for optimal separation. Separated peptides were sprayed into an Orbitrap Astral mass spectrometer at an electrospray voltage of 3.5 kV. Precursor ions ranging from 380-980 m/z were measured in the Orbitrap analyzer at a resolving power of 120,000, requiring either 1,000,000 charges or a 50 ms maximum injection time for each scan. Precursors from 380-980 m/z were selected for MS2 using DIA and 4 Th quadrupole isolation windows (150 total windows across the mass range). Normalized collision energy (NCE) was set to 27%. Fragment ions from m/z 100-1,000 were measured in the Astral analyzer, requiring either 50,000 charges or a 3.5 ms maximum injection time.

**Figure 1.**
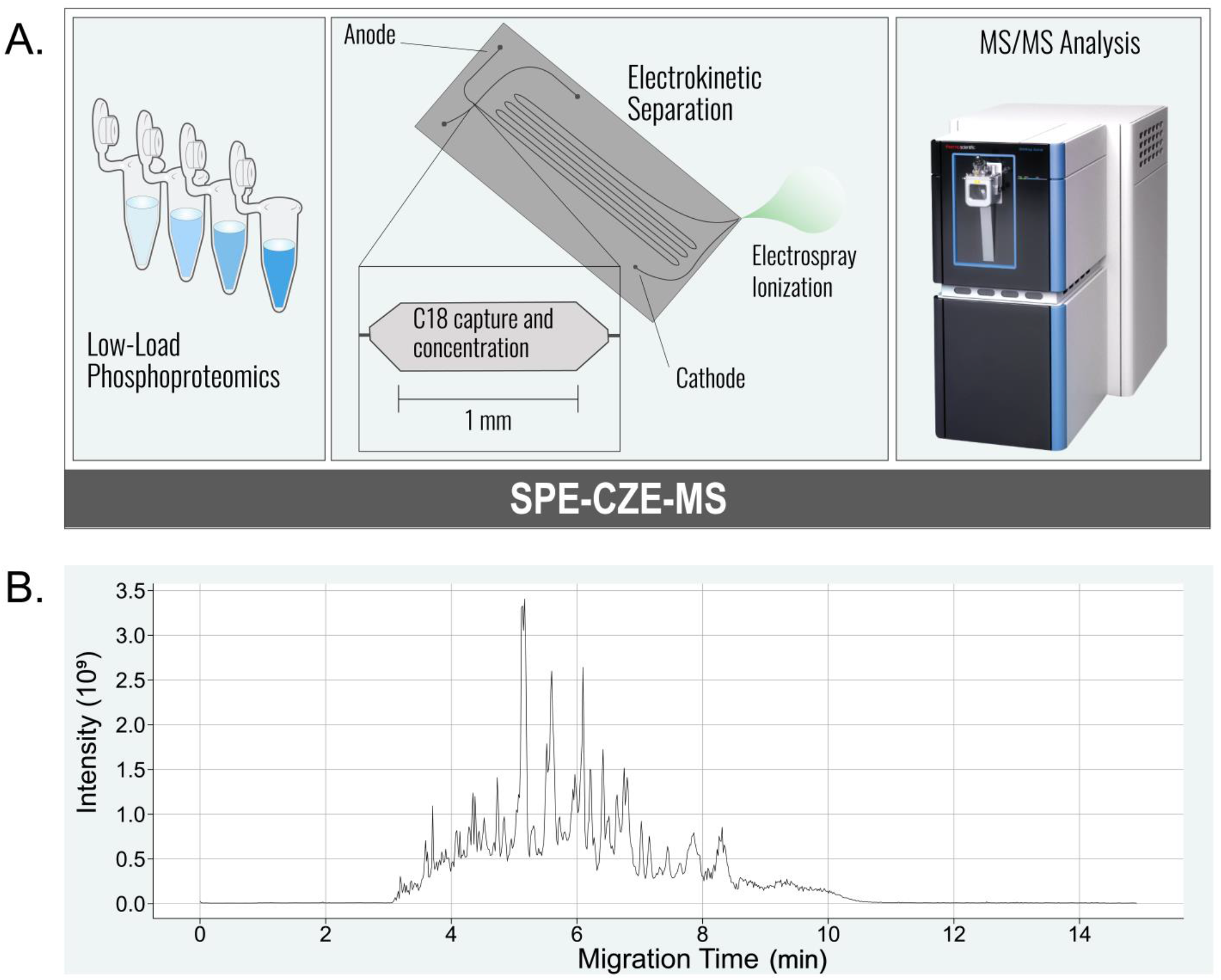
SPE-CZE-MS workflow for low-load phosphoproteomics. **(A)** Five-fold serially diluted synthetic phosphorylated peptide standards or phosphorylated peptide-enriched mouse brain tissue were loaded onto the SPE-CZE device and electrokinetically separated over 15 min using a 500 V/cm electric potential before Orbitrap Astral DIA analysis. **(B)** Electropherogram of 32 ng on-chip analysis of enriched mouse brain tissue showing an 8-minute separation window.

### Data analysis

To analyze the synthetic standards collected on either SPE-CZE-MS or nLC-MS. Thermo Scientific .RAW files were uploaded to Skyline (version 23.1).^47,48^ Extracted ion chromatograms (XIC) were mined for all doubly, triply, and quadruply charged precursors of all targets, rendering a final set of 392 precursors. Only singly charged fragment ions were extracted. Mass tolerances for MS1 and MS2 were set to 10 ppm. XICs of both precursor and ions were manually validated. Linear regression was performed on the log_10_ -transformed integrated peak areas (summed area of all transitions) exported from Skyline as a function of log_10_-transformed concentrations using the “linregress” function from the SciPy python package (v. 1.10.0) stats module.

For phosphorylated peptide-enriched sample analysis, Thermo Scientific .raw files collected using SPE-CZE-MS or nLC-MS were searched separately on Spectronaut (Biognosys, version 19.3.241023). For the initial database search used to generate a predicted DIA library, the files were searched with the Pulsar engine against a FASTA file containing 63,755 sequences from the Swiss-Prot UP000000589 *Mus musculus* database downloaded on 03/23/2021. Tryptic peptides between seven and 52 amino acids with at most two missed cleavages were considered. Carbamidomethylated cysteine was set as a fixed modification and phosphorylated serine, tyrosine, and threonine, acetylated protein N-term, and oxidized methionine were set as variable modifications—allowing up to five per peptide. The first search to determine systematic mass error was set to 40 ppm before implementing a narrower, dynamic search tolerance for peptides to be included in the directDIA library. For the predicted library search, the decoy generation method was set to “Mutated”. Protein identifications were filtered to meet a 1% protein-level false discovery rate (FDR) threshold and the IDPicker^49^ algorithm was used for protein inference. For SPE-CZE-MS, all retention time calibration settings were disabled. All reported phosphorylated peptides have at least a 0.75 localization probability. Other settings not noted here were set to default.

### Variable definitions and equations

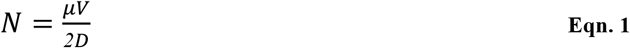

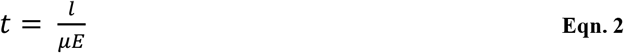

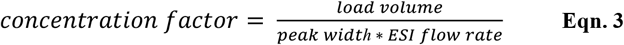

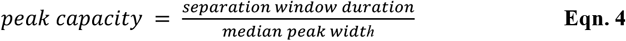

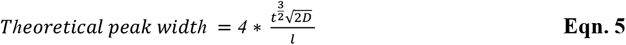

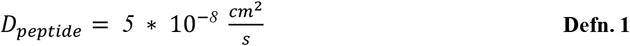

where *N* is the number of theoretical plates, µ is the electrophoretic mobility, *V* is the applied voltage across the channel, *D* is the solute diffusion coefficient, t is the migration time, *l* is the length of the separation channel, and *E* is the electric field strength (*V* / *l*).

## RESULTS AND DISCUSSION

### Design and rationale

CZE has proven to be a viable separation method for complex mixtures due to its successful coupling to electrospray ionization and its potential to deliver very fast yet high-resolution separations. To fully take advantage of the fast and high-resolution abilities of CZE-based separation for complex mixtures, many advancements toward interfacing it with electrospray ionization as well as integrating pre-concentration techniques have been pivotal. The CZE device described here leverages an SPE-based pre-concentration method and an integrated ESI emitter (**Figure 1A**). This study aims to quantitatively validate this analytical approach for MS-based single-cell equivalent phosphoproteomics. Rapid separations achieved with minimal sample loss at transfer steps is crucial in pushing the sensitivity limits for MS/MS analysis of single-single cell -omics. Here, we characterize separation and quantitative performance for 225 phosphorylated peptide standards from 2 fmol down to 5 zmol on-chip as well as phosphorylated peptide-enriched mouse brain tissue from 32 ng (**Figure 1B**) down to 50 pg loaded on-chip. Our goal is to define important analytical limitations of the SPE component within the chip-based CZE device for phosphorylated peptides, assess the compatibility of our separation method with the Orbitrap Astral, and validate quantitative performance of complex phosphoproteomics mixtures.

### Load volume characterization

Tryptic digests of whole proteome samples produce peptides with a range of physicochemical properties that are relevant to the retention of C18 solid-phase material. The presence of a phosphoryl group can render hydrophobic-based retention a less favorable chemical interaction by increasing the polarity of the analyte. Tenuous retention of a relatively more hydrophilic subset of peptides can be impacted by solvent volume overload on the 5.7 nL SPE bed. We tested these limitations on a set of 225 phosphorylated peptide standards ranging in hydrophobicity and number of phosphorylation modifications. Standards were reconstituted and diluted to a concentration of 200 amol/μL and loaded onto the SPE bed for analysis for 30 sec, 1 min, 2 min, 4 min, or 8 min. The flow rate of the loading is approximately 500 nL/min, rendering the loading volumes 0.25 μL, 0.5 μL, 1 μL, 2 μL, or 4 μL for each condition, resulting in loading amounts of 50 amol, 100 amol, 200 amol, 400 amol, and 800 amol, respectively. Two technical replicates were collected for each condition. The peak area should increase as we increase loading amount on the SPE bed if mass or volume overload does not occur. Mass overload can occur if there is not enough solid phase material to retain the amount of sample. Too much volume flowing through the bed can impact subpopulations of peptides that form relatively weaker interactions with the solid phase material, such that the forces of the fluid flow are strong enough to overcome the hydrophobic interaction of weakly retained analytes through the bed.

For the loading volume experiments, we used an optimized SPE-CZE-MS DIA method for analyte detection and quantification. Integrated phosphorylated peptide peak areas were calculated and exported from Skyline after manual inspection of peaks. The peptides were binned by nLC-MS retention time as a proxy for hydrophobicity and the median intensity that is normalized to the highest intensity is plotted as a function of loading amount (**Figure 2D**). Several trends highlighted here provide possible mechanistic explanations for volume overload. First, on a universal level, the signal does not increase linearly despite loading more material, albeit across higher volumes. This was not the case when controlling for volume (4-min loading time maintained) (**Figure 2E**). The rate and direction of change in intensity from 4 to 8 min also provides interesting information about how each of these subpopulations could be interacting with the SPE bed. The most hydrophilic subset (early eluting analytes on nLC) shows the lowest amount of material retained, evidenced in its apex at the 2 min loading time with the lowest median intensity. The relatively sharp decrease in signal between 2 and 4 min suggests that this subpopulation of peptides is retained well before a volume overload at 4 min. The consistent low signals at 4 and 8 min suggests most of these peptides flowed through before the intended elution. The next most hydrophilic set shows higher apexes at both 2 and 4 min and complete dissociation at 8 min. It is possible that some of these peptides that apexed at 2 min were out-competed by a more hydrophobic set during the 4-min load, preventing the association of more material from this group onto the SPE bed. The two next most hydrophilic sets show an even higher median intensity apex at 4 min and no change or very minimal decrease at 8 min, suggesting a similar effect where more hydrophobic peptides are preventing the retention of additional material from this set. Finally, the most hydrophobic set of peptides show the highest median-normalized intensity at 4 min and appear to drop in intensity most steeply from 4 to 8 min, suggesting these peptides were well-retained until a point of volume overload at 8 min. A possible remedy for this effect of volume overload is a larger SPE bed to prevent sample loss during this process. The percent coefficient of variation distributions at each load time indicate improved reproducibility for all hydrophobicity bins at 4 min relative to 8 min (**Figure 2C, Supplementary Figure S3**). These results directed our decision to load for 4 min for all subsequent analyses.

**Figure 2.**
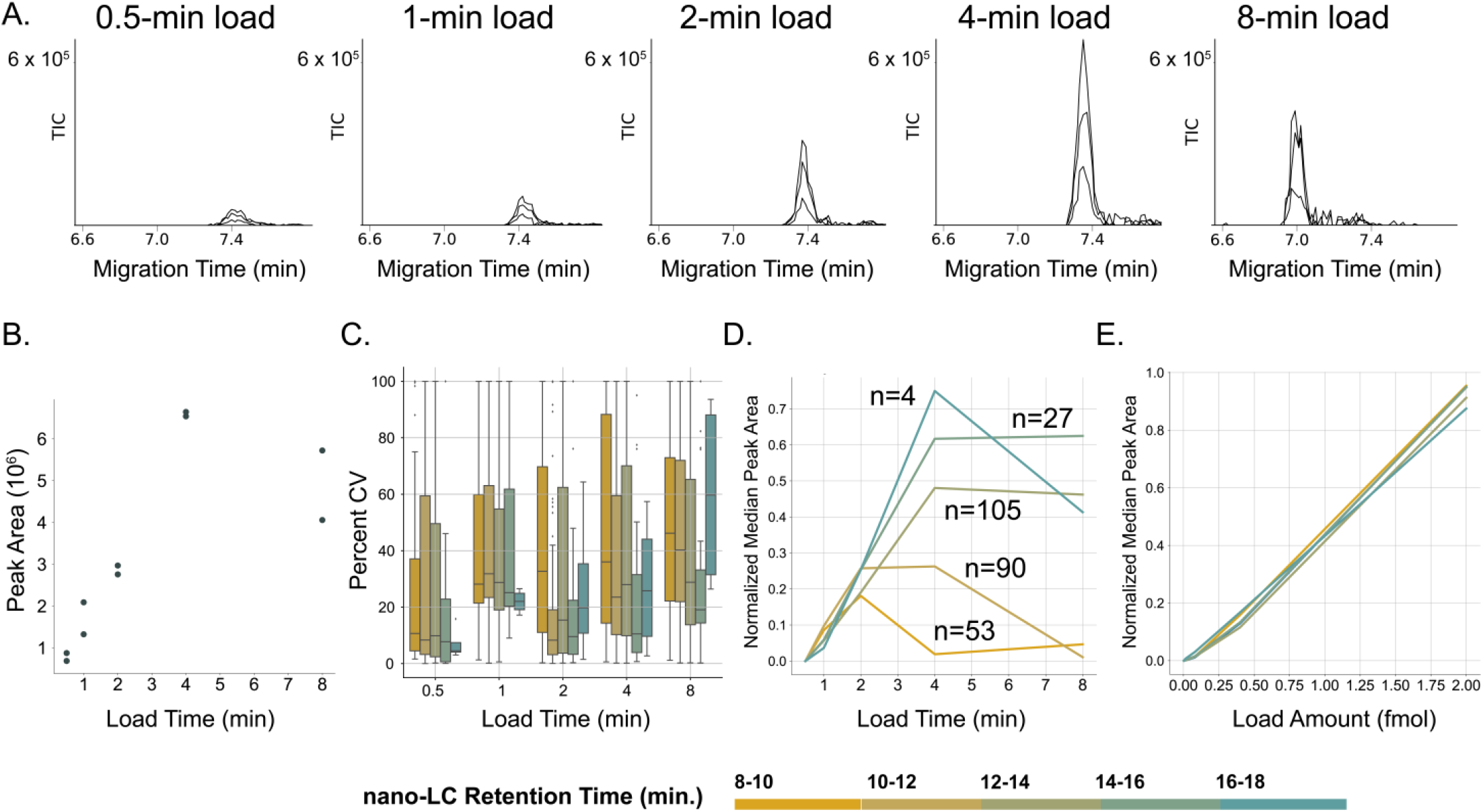
Volume overload characterization. **(A)** Electrophoretic peaks for peptide GT(phos)PPLTPSDSPQTR ++ for loading times 0.5, 1, 2, 4, 8 min that apexes at 4 min. **(B)** GT(phos)PPLTPSDSPQTR ++ integrated peak area as a function of load time. **(C)** Percent CV distributions of standards for each load time. **(D)** Manually inspected precursor ions (n=558) were binned into nLC retention time bins as a proxy for hydrophobicity measures. Each set were normalized to the highest intensity value per response curve. The median of each bin plotted as a function of load time. **(E)** The same precursor peak areas are plotted as a function of loading amount with loading time of 4-min was held constant.

### Separation performance of SPE-CZE device

We next characterized the separation performance and concentration factors using synthetic standards. As expected, we see an increase in the full width-half maximum (FWHM) of the electrophoretic peaks as a function of migration time (**Eqn. 2**) (**Figure 3A, B**). Note that the FWHM is based upon the mean value of all considered transitions for each standard as calculated by Skyline. In line with expectation, the concentration factors decrease with increasing peak width (**Figure B, Eqn. 3**).

**Figure 3.**
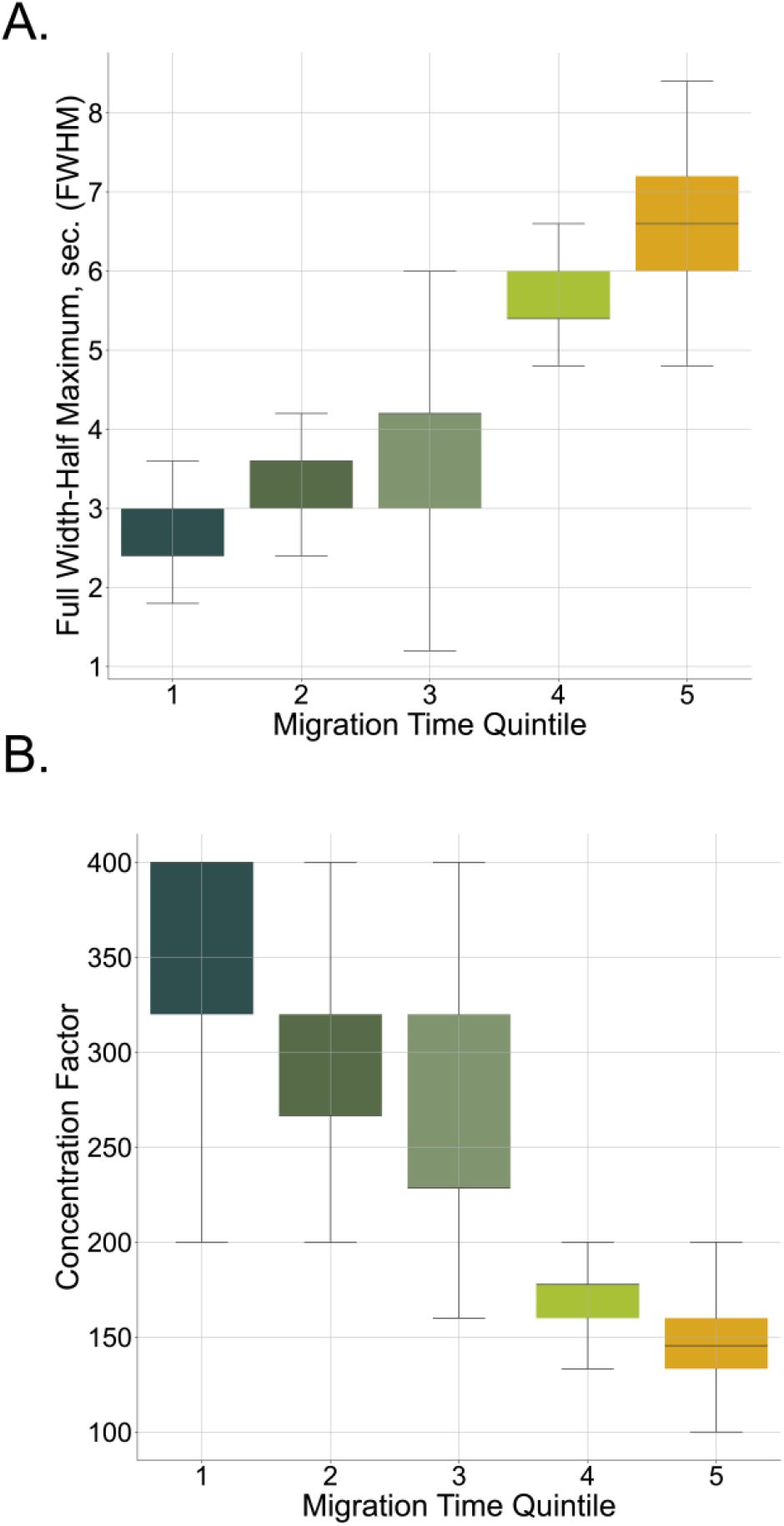
Separation characterization for synthetic standards and complex mixtures. Full width-half maximum values for 2 fmol standards **(A)** and 32 ng load enriched mouse brain tissue **(B)** analyzed with SPE-CZE-MS binned by migration time quintile. Concentration factor distributions of 2 fmol standards **(C)** and 32 ng load enriched mouse brain tissue **(D)** binned by migration time quintile.

### Benchmarking quantitative performance SPE-CZE analysis of synthetic standards to nLC-MS

We next analyzed the signal response of each standard to 5-fold dilutions from 2 fmol to 1 zmol when analyzed with SPE-CZE-MS and compared results to the same samples analyzed with nLC-MS. Due to the highly disparate technological features and analytical conditions of SPE-CZE and nLC-MS, we decided to analyze our dilution curve using the best-performing mass spectrometry method independently designed for each device, yet maintaining the active separation time constant. We injected the same loading amount for analysis on either SPE-CZE-MS or nLC-MS.^50^ For each SPE-CZE injection, the sample was loaded for 4 min (∼2 μL) onto the SPE bed after the bed was conditioned with weak mobile phase. Phosphorylated peptides were then eluted from the bed using BGE and positive pressure applied to the sample well. Additional positive pressure was applied to the cathode to facilitate sample transfer into the 22 cm separation channel for electrokinetic focusing and separation. The 500 V/cm electric field was maintained constantly for 15 min and the separation window for each run was approximately 8 min. After electrokinetic separation, peptides were sprayed into an Orbitrap Astral mass spectrometer at an electrospray voltage of +3.5 kV and analyzed using an optimized data independent acquisition (DIA) method (**Methods, SPE-CZE-MS)**. Phosphorylated peptides analyzed using nLC-MS were chromatographically separated using a 30 cm nano-capillary column^46^ at a flow-rate of 300 nL/min over a 16-min active gradient, implementing a 4-min wash step and 16-min re-equilibration step. Peptides were sprayed into the Orbitrap Astral mass spectrometer at an electrospray ionization voltage of 2 kV and collected using a DIA method developed elsewhere (**Methods, nLC-MS**).^48^ Two technical replicates were acquired for each sample. Note that among several different parameters, the SPE-CZE method utilizes 4 Th bins unlike the 2 Th bins implemented in the nLC-MS method. We acknowledge that different parameters may be ideal for quantification than those described in this study that we optimized for sensitivity. Here, we investigate how each respective method performs quantitatively across our dilution series of synthetic standards. We plotted the coefficients of determinations for all standards (including charge states 2-4) for a 3-point curve down to a 10-point curve (full set from 2 fmol to 1 zmol). **Figure 4A** shows these calculations for SPE-CZE-MS-quantified standards (top) and nLC-MS-quantified standards (bottom). The shaded regions highlight standards quantified with coefficients of determination > 0.9. Venn diagrams above each curve (3-point, etc.) show the overlap of these subsets of standards (R^2^ > 0.9) between each respective separation method, wherein dark blue codes for SPE-CZE-MS, light blue codes for nLC-MS, and shaded yellow codes for overlap of the two methods. 60.0% percent of SPE-CZE-quantified standards are linear down to 640 zmol. Only 31% percent of these were determined to have coefficients of variation under 20% (median CV= 40.1%). These values for nLC-MS were 17.4% percent down to 640 zmol with 12% coefficients of variation (CV) under 20% (median CV = 45%) (**Figure 4C**). The greater number of points per peak (FWHM) for SPE-CZE because of wider DIA windows (**Figure 4B**) could explain the enhanced quantification achieved by SPE-CZE-MS. Note that this plot does not include MS1 scans which are shown in Supplementary Figure S4. Despite the enhanced linearity of the SPE-CZE method, the nLC-MS quantitative reproducibility for peptides quantified with at least a 0.9 coefficient of determination comparable to SPE-CZE-MS down to 128 zmol concentrations, which may suggest that the quantification may have not been severely compromised from the narrower windows. From these results, we conclude that SPE-CZE-MS can reliably generate a response for ∼60% of peptides commensurate with loading down to 640 zmol and spanning 5 orders of magnitude.

**Figure 4.**
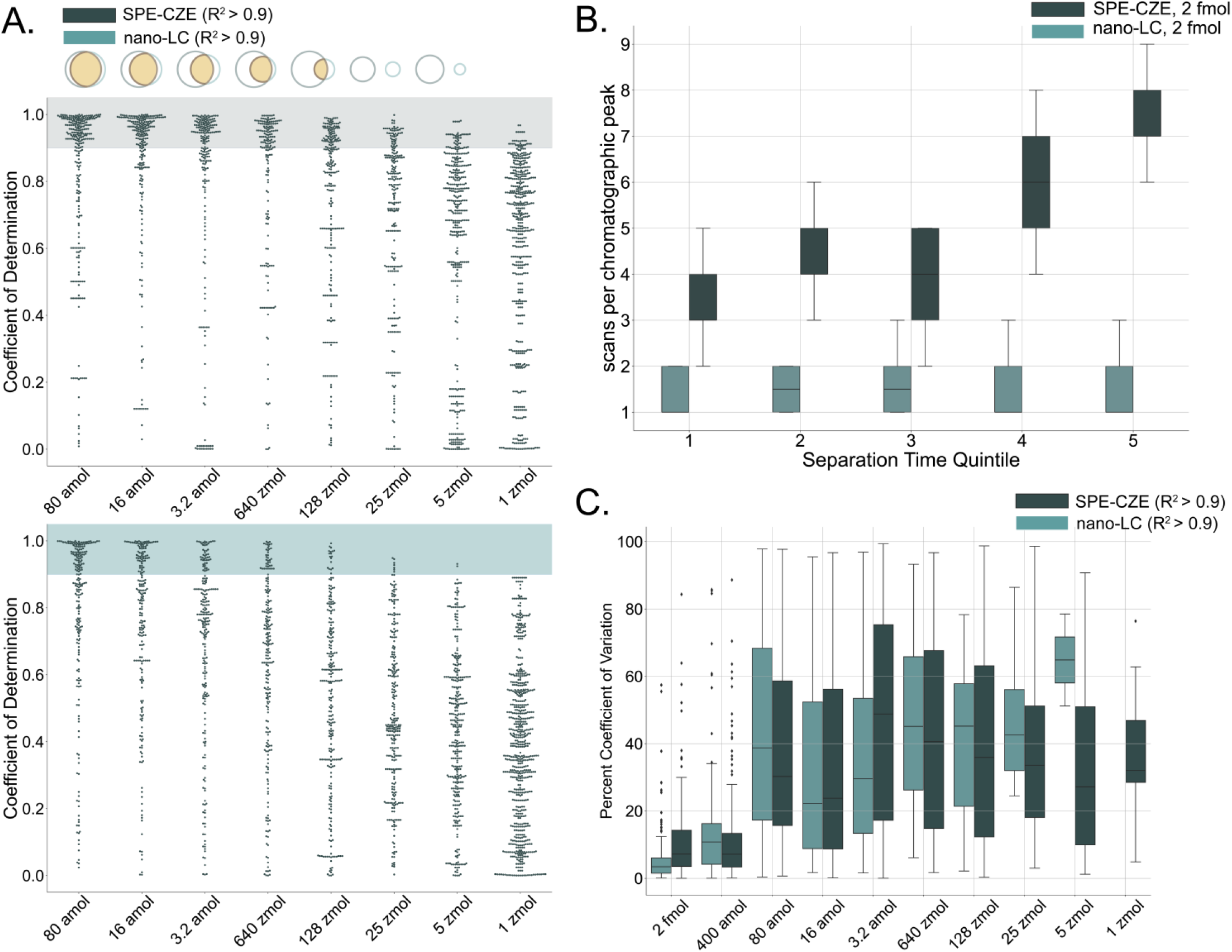
Quantitative performance assessment with synthetic phosphorylated peptide standards. (A) Phosphorylated peptide standard response curves were generated for 3-point curves (down to 80 amol), 4-point (16 amol), 5-point (3.2 amol), 6-point (640 zmol), 7-point (128 zmol), 8-point (25 amol), 9-point (5 zmol), and 10-point (1 zmol) for samples analyzed with either SPE-CZE-MS or nLC-MS . Coefficients of determination are plotted for each curve for SPE-CZE-MS (top) and nLC-MS (bottom). **(B)** Distributions of MS2 scans per electrophoretic peak (FWHM) as a function of separation time quintile for each separation method for the 2 fmol loading amount. **(C)** Distributions of percents coefficient of variation for replicates of standards that had response curves with r-squared values >0.9.

### Quantitative performance of decreasing loads of phosphorylated peptide-enriched sample

We next analyzed the phosphorylated peptide-enriched mouse brain tissue data using Spectronaut, filtering for 1% peptide and protein FDR, and requiring at least 0.75 phosphorylation site localization probabilities. The number of phosphorylated peptides, phosphorylation sites, and phosphorylated proteins were comparable across all loading amounts for each respective separation method, except for the 32-ng load (**Figure 5A**). Each 5-fold dilution exhibits roughly a five times difference in identifications except for the 32 ng to 6 ng dilution. Possible reasons for this are being explored to improve dynamic range at the upper limit for SPE-CZE. All identified peptides migrated within 9.3 min. The median peak width of 5.04 s therefore rendered an approximate peak capacity of 110.6 (**Eqn. 4**) These specifications of the complex mixture very closely match the theoretical calculations plotted in **Supplementary Figure S1**.

**Figure 5.**
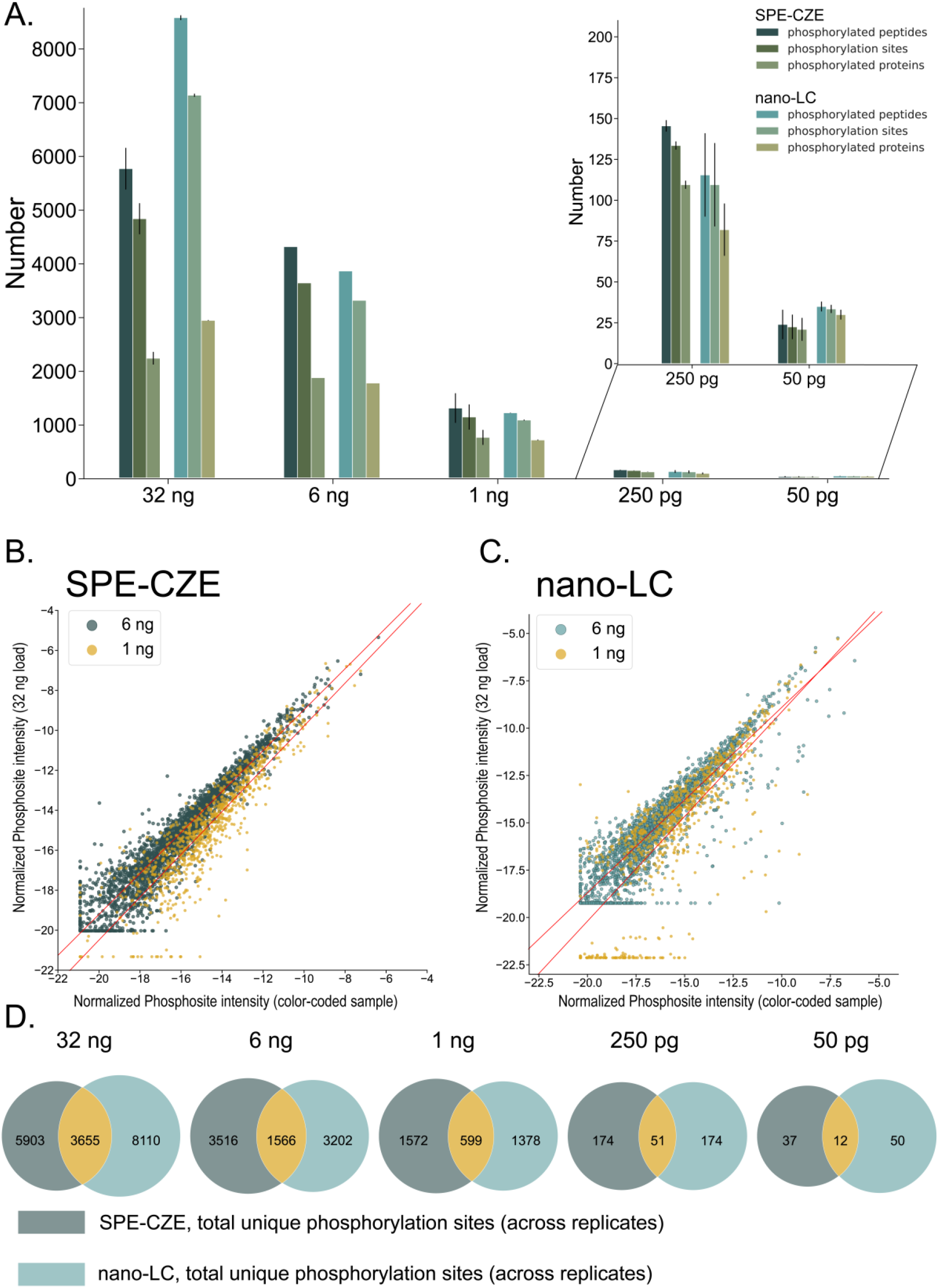
Characterization and quantitative performance assessment for enriched phosphorylated peptides. (A) Average phosphorylated peptides, sites and proteins for each loading amount of phosphorylated peptide-enriched mouse brain tissue analyzed on SPE-CZE-MS or nLC-MS. Phosphorylation site intensities relative to the summed intensities for the 32-ng loaded sample on SPE-CZE-MS **(B)** and nLC-MS **(C)** are plotted against that for the 6 ng and 1 ng samples. Regression analysis determined r-squared values of 0.95 and 0.9 for 6 ng and 1 ng samples analyzed on SPE-CZE-MS respectively, and 0.91 and 0.79 for the 6 ng and 1 ng analyzed on nLC-MS, respectively. The standard errors for these best fit lines on SPE-CZE-MS are 0.006 and 0.018 while that for nLC-MS are 0.009 and 0.031. **(D)** Overlap of phosphorylation sites across loading amounts from 32 ng to 50 pg.

We then tested how well the relative quantification was maintained from the highest loading amount (32 ng) down to 1 ng for SPE-CZE-MS (**Figure 5B**) and nLC-MS (**Figure 5C**). We normalized each protein in the 32-ng sample loading amount to the summed protein intensity from that sample. The same calculation was made for the 6 ng and 1 ng loading amounts and are plotted in green and yellow, respectively. Replicates were averaged for each condition. We plotted the best-fit line derived from linear regression analysis and observe correlation values of 0.95 and 0.9 for the 6 ng and 1 ng measured with SPE-CZE-MS and 0.91 and 0.79 for that measured with nLC-MS. Encouragingly, the standard error for the SPE-CZE-MS best-fit lines were 0.006 and 0.018. These results suggest that relative quantification is reproducible when scaling down sample loading to 1 ng on-chip. Finally, we compare the phosphorylation sites returned by SPE-CZE-MS and nLC-MS and observe a high degree of complementarity (**Figure 5D**). This implies that both methods are useful for discovery, perhaps in tandem, by enabling the measurement of phosphorylation sites harbored on peptides ranging in physicochemical properties.

### Distinct selectivity for complex phosphoproteomics mixtures between SPE-CZE-MS and nLC-MS

The physicochemical enrichments of certain phosphorylated peptides in SPE-CZE did not suffice to explain the high degree of complementarity between the two separation methods (**Supplementary Figure S5**). We figured the differing results at the phosphorylation site level were largely driven by the different selectivity afforded by the complementary separation mechanisms (**Figure 6A**). One particularly relevant aspect of the separation mechanism of SPE-CZE to the application of phosphoproteomics is shown in **Figure 6B**. Phosphoryl groups have two sites that must be protonated to neutralize that moiety on the peptide. It is likely therefore that some sub-population of the phosphorylated peptides at pH 2 will have lower charge states than would peptides without that additional proton-accepting moiety, resulting in a slower velocity. We observe a global separation of phosphorylated peptides from non-phosphorylated peptides from SPE-CZE analysis (**Figure 6B**, top), whereas the separation mechanism of nLC-MS does not result in this effect (**Figure 6B**, bottom). We find that this confers a benefit on the analysis of multiply phosphorylated peptides. Firstly, despite the greater sensitivity afforded by nLC-MS at 32 ng loading, SPE-CZE detected a greater number of multiply phosphorylated species. These multiply-phosphorylated peptides were detected starting at five min– the point in the separation window at which the median number of non-phosphorylated peptide analysis drops steeply. The multiply phosphorylated peptides detected in the nLC-MS separation elute consistently across the active separation gradient (**Figure 6C**). We then compared all the localization probabilities of each phosphorylation site on multiply phosphorylated peptides and plotted localization probability against ranked localization probability for SPE-CZE-MS and nLC-MS (**Figure 6D)**. Note, we show phosphorylation sites with probabilities below 0.75 for this analysis. We observe that not only are we detecting a higher number of multiply phosphorylated peptides in the SPE-CZE-MS analysis, but also producing spectra for these species that enable higher-confident localization. This feature specific to the separation mechanism of SPE-CZE is particularly beneficial for our 4 Th DIA windows, as it may serve as compensation for the relatively lower quadrupole selectivity compared to the 2 Th method many have validated for nLC-MS.

**Figure 6.**
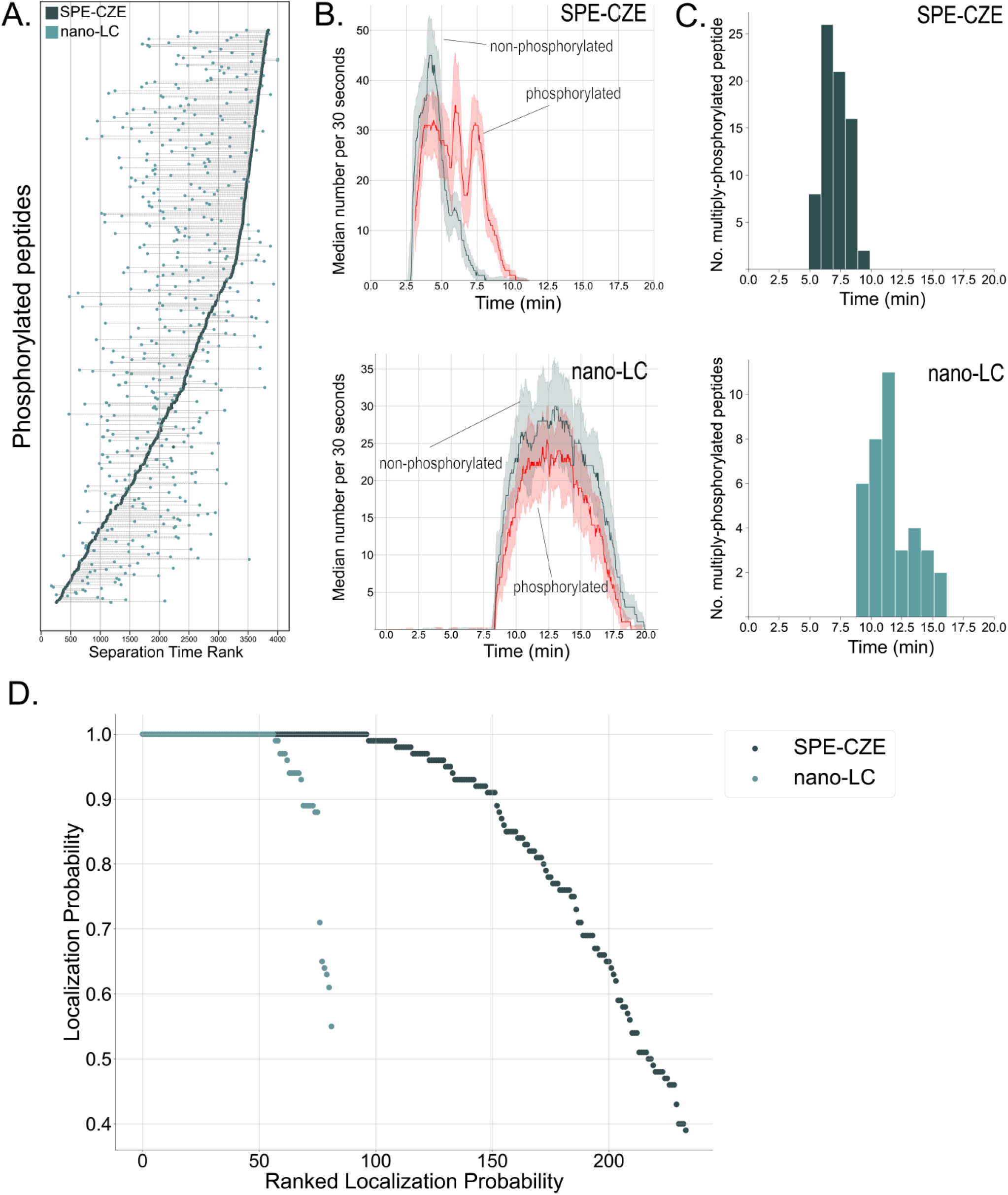
Complementary selectivity between SPE-CZE-MS and nLC-MS confers benefits for multiply phosphorylated peptide analysis. **(A)** Ranked separation time from SPE-CZE-MS and nLC-MS analysis are plotted for each shared phosphorylated peptide from the 1 ng load. **(B)** The median and standard deviation of number of phosphorylated (red) or non-phosphorylated peptide (dark blue) identification (binned in increments of 30 seconds) over separation time for SPE-CZE-MS (top) and nLC-MS (bottom). **(C)** Number of multiply phosphorylated peptides over migration time window for SPE-CZE-MS (top) and nLC-MS (bottom). **(D)** Localization probability as a function of ranked localization probability for multiply phosphorylated peptides analyzed with SPE-CZE-MS or nLC-MS.

## CONCLUSION

The integration of the SPE bed in the microchip-based CZE device used in this study attempts to incorporate the pre-concentration benefits inherent to chromatographic separations while leveraging electrokinetic-based separations. While this device has been demonstrated to enhance the sensitivity for proteomics analysis, we assessed its performance for MS-based phosphoproteomics analyses, which suffer from even greater sample limitations. These analytes exhibit globally divergent physicochemical characteristics from unmodified peptides, thus requiring experimentation to establish optimal methodological parameters for use with the SPE-CZE device, and perhaps differing architectural needs such as the size of the SPE-CZE bed. We show that the current dimensions of the SPE bed can retain sample amounts up to at least 2 femtomolar. Experiments where we load this amount but with a higher volume than 2 μL, we observe symptoms of volume overload which will be considered in future designs of the device.

This study also explores the compatibility of the Orbitrap Astral mass spectrometer with the electrophoretic timescale of 8-minute electrokinetic separations. The electrophoretic peak width proportionally narrows with a decrease in separation time. Thus, both detection and extracted-ion-electropherogram-based quantification can be impacted if the mass spectrometer scan speed is too slow for sufficient peak sampling. Here, through analyzing 225 phosphorylated peptide standards with SPE-CZE-MS, we show that the median scans per peak throughout the migration time window ranges from 4 to 9. The same standards were able to be quantified with high linearity (>0.9) down to 640 zmol. The reproducibility of the measurements with high linearity were highly reproducible down to 400 amol with percents coefficient of variation below 15% but increased to 20-40% for lower concentrations. A similar trend was observed with nLC-MS measurements. Future studies will attempt to investigate this effect further, but one possibility is the tenuous retention of phosphorylated peptides on hydrophobic surfaces, resulting in inconsistent sample concentration on the SPE bed. When expanding our quantitative benchmarking to phosphorylated peptide-enriched mouse brain tissue, we show that the relative quantification of phosphorylated peptides is maintained with decreasing sample amount. Finally, we explore how the differing separation mechanisms between CZE and nLC confer distinct selectivities that result in complementary detection of phosphorylation sites. We find that this effect benefits the detection and localization confidence of multiply phosphorylated peptides with SPE-CZE analysis. Our findings support that SPE-CZE-MS is a sensitive method for quantitative analysis of low-load phosphoproteomics and can confer a selectivity benefit for multiply phosphorylated peptides.

## DATA AVAILABILITY

The raw data for proteomics datasets in this study have been deposited to the MassIVE database under the accession number **MSV000096595**

Username: **MSV000096595_reviewer**

Password: 5TzHLKlM1NDCj9Hm

## ACKNOWLEDGEMENT

This work was supported in part by a sponsored research agreement with Thermo Fisher Scientific (JJC), the National Institute of General Medical Sciences of the National Institutes of Health (grants P41GM108538 and R35GM118110 to JJC), and the National Human Genome Research Institution through a training grant to the Genomic Science Training Program (grant T32HG002760 to LRS). We also acknowledge support from US DOE Award Number DE-SC0018409. The content is solely the responsibility of the authors and does not necessarily represent the official views of the National Institutes of Health.

## AUTHOR CONTRIBUTIONS

LRS, JSM, JWT, and JJC Conceptualization; LRS, JSM, JWT methodology, LRS and JSM Validation; LRS and JSM Formal Analysis; LRS, NML, MLR, JSM, and JWT Investigation; JSM, JWT, and JJC Resources; LRS Data Curation; LRS Writing-Original Draft; LRS, JSM, KAO, STQ and JJC Writing-Review & Editing; LRS, JSM, JWT, KAO, STQ Visualization; JSM, JWT, and JJC Supervision; JSM, JWT, and JJC Project Administration; JJC Funding acquisition

