## Supplementary Figures for "SPE-CZE-MS quantifies zeptomole concentrations of phosphorylated peptides"

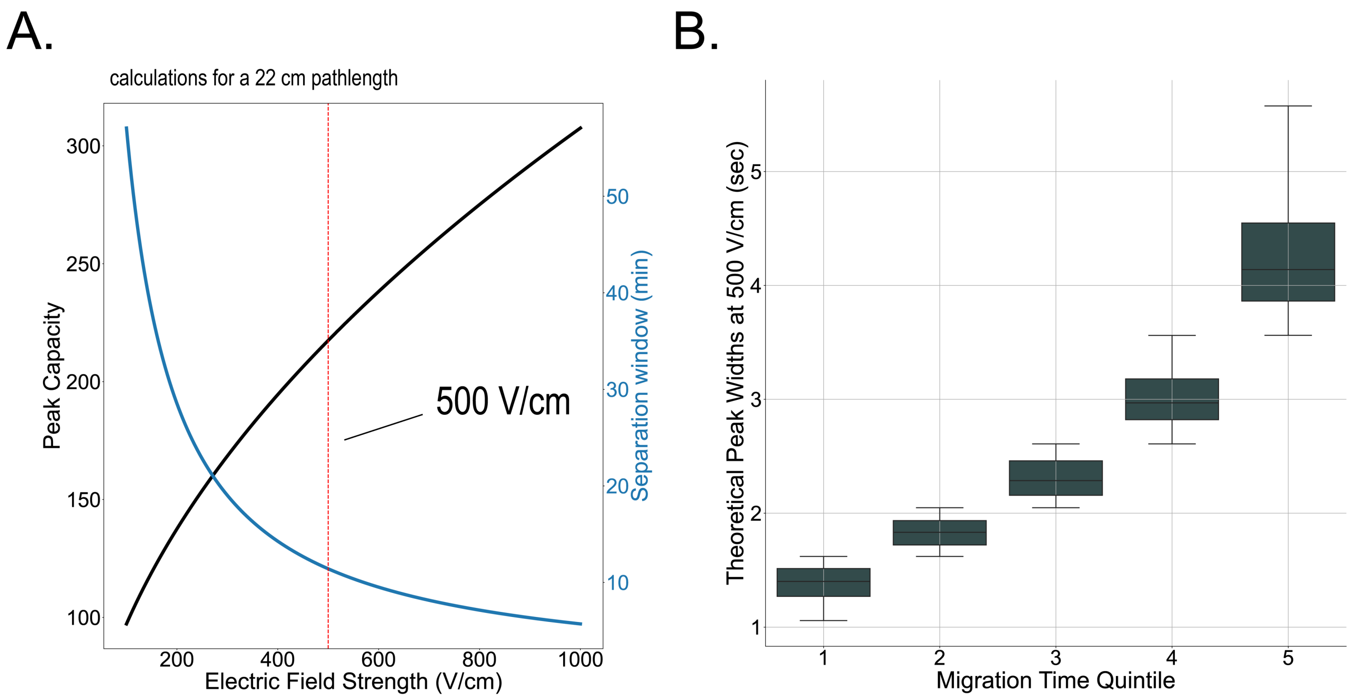


**Supplementary Figure S1. (A)** Theoretical peak capacity and total separation duration as a function of electric field strength. A red dotted line indicates 500 V/cm. **(B)** Theoretical peak widths (**Eqn. 5**) at 500 V/cm for 5 sections of the migration window.

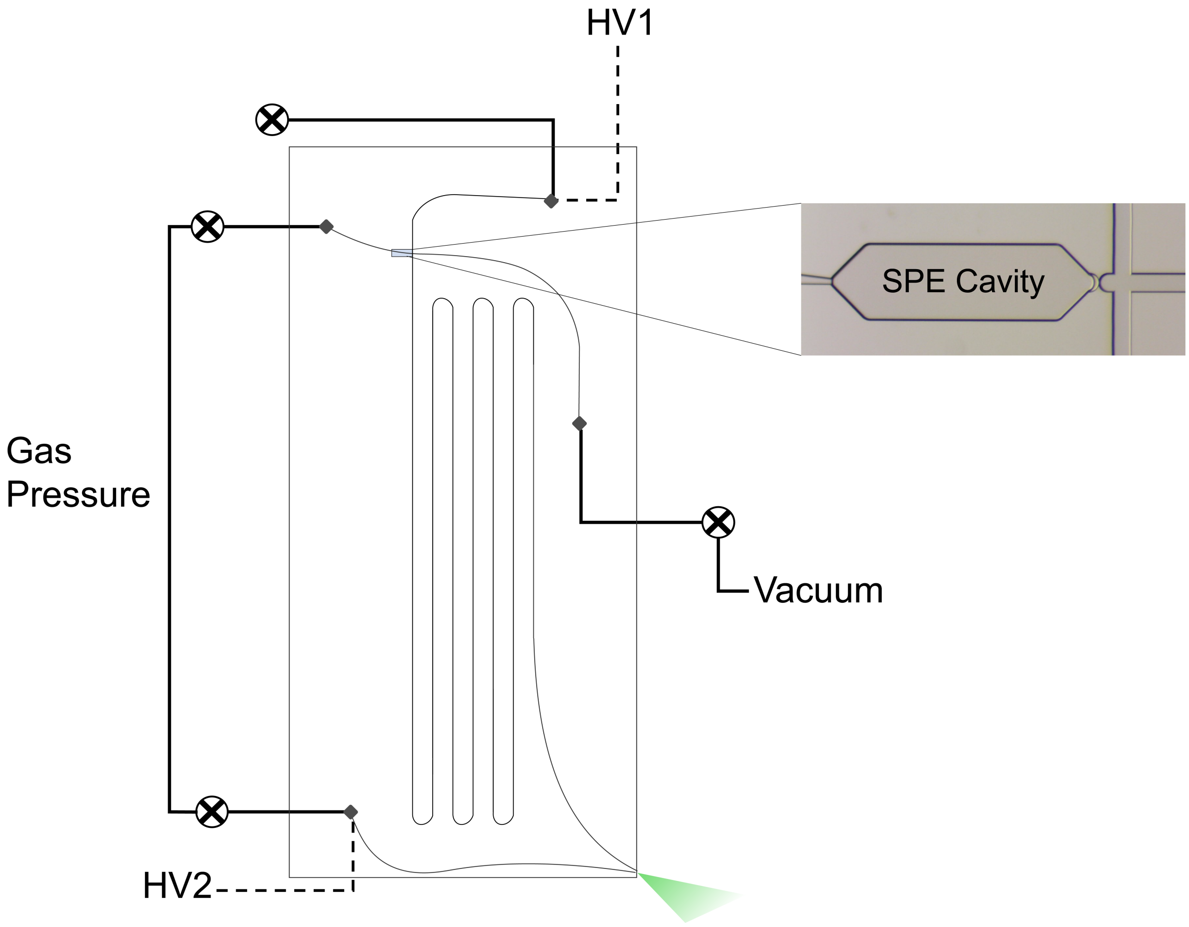


**Supplementary Figure S2. SPE-CZE diagram.** Positive pressure applied to sample inlet and electrode. Vacuum applied to waste. HV1 is set to 15 kV and HV2 is set to 2.4 kV to achieve a 500 V/cm electric field and a 3.5 kV electrospray ionization voltage at the corner of the chip. The 1 mm length SPE bed is situated downstream of the sample inlet.

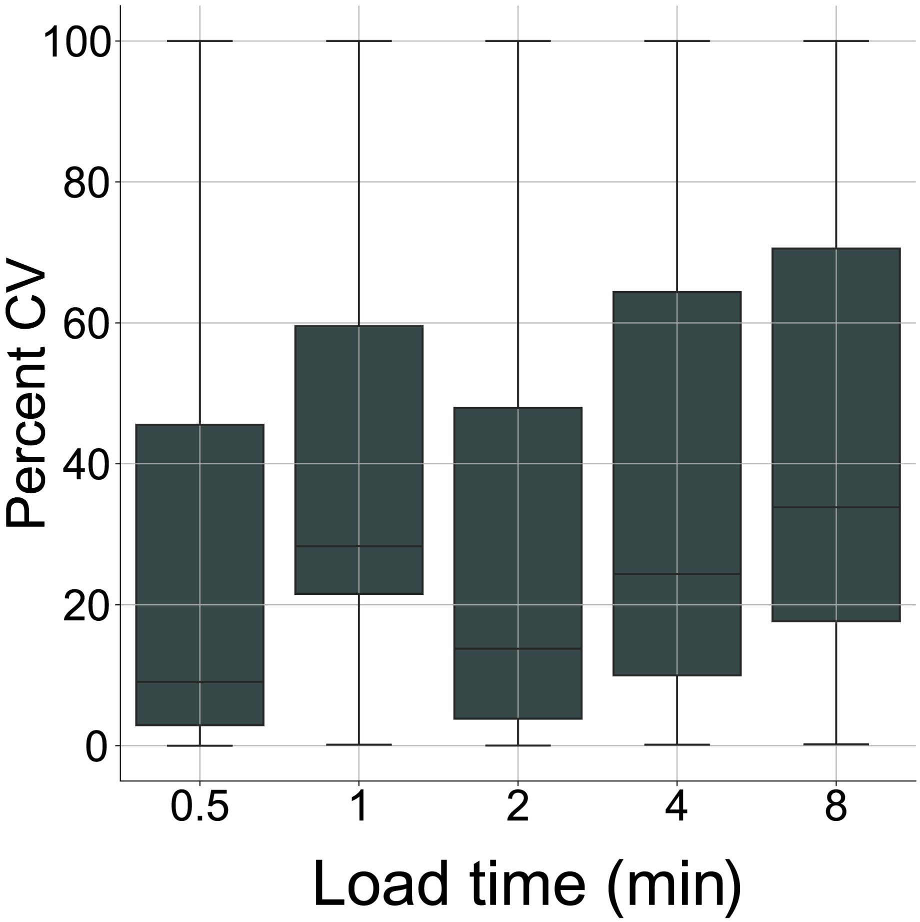


**Supplementary Figure S3. Percent coefficient of variation distributions of synthetic phosphorylated peptide intensities for each load time tested on SPE-CZE-MS.** Colors code for time window during which the peptides eluted in the described nLC-MS experiment as a proxy for hydrophobicity.

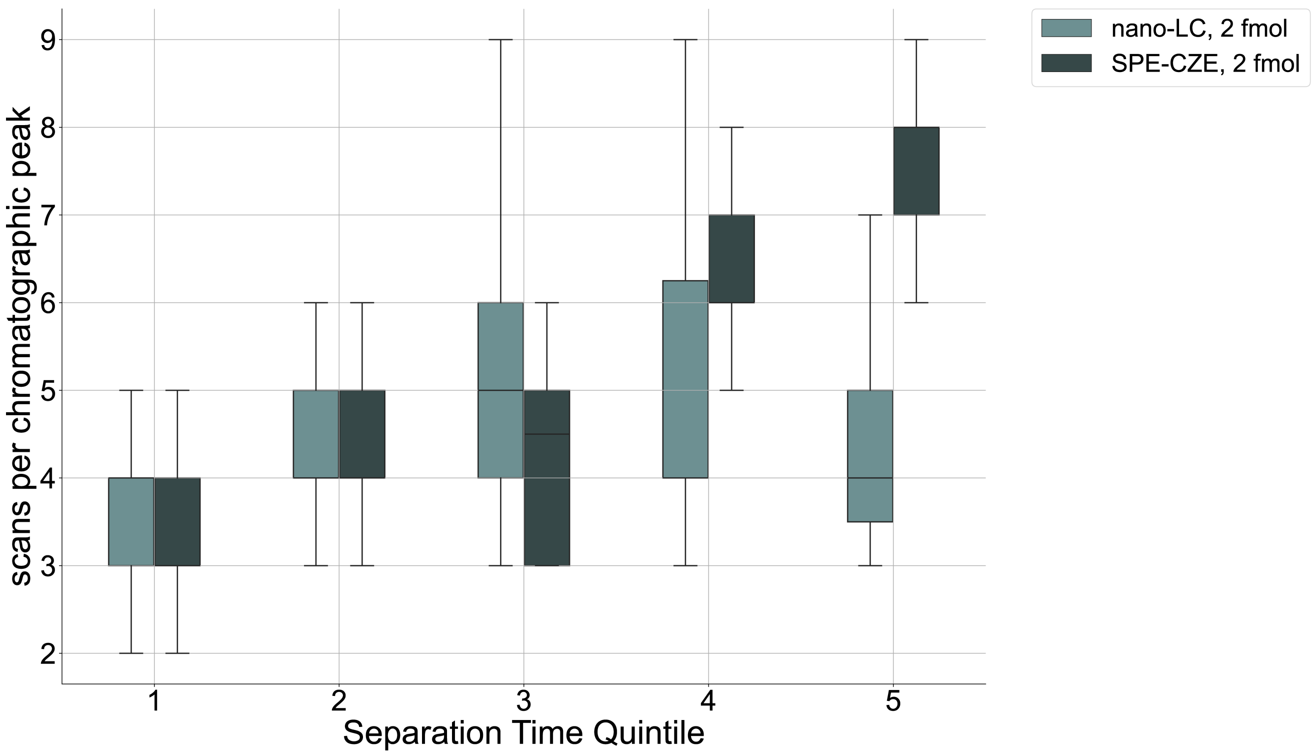


**Supplementary Figure S4. MS1 scan distribution per peak across the separation time range for nLC and SPE-CZE.**

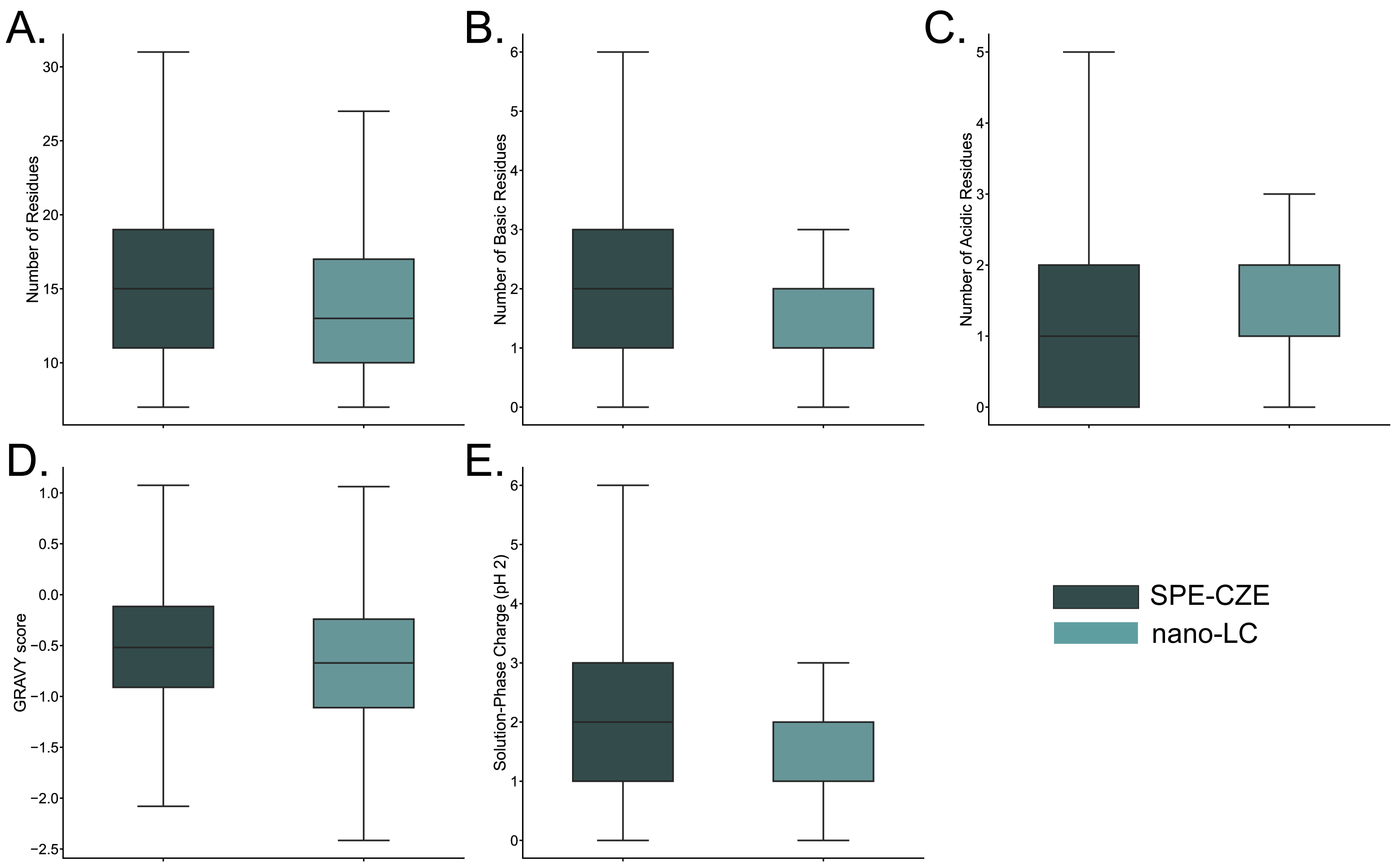


**Supplementary Figure S5. physicochemical enrichments in SPE-CZE vs. nLC peptides including length (A), number of basic residues (B), number of acidic residues (C), GRAVY score (D), and solution-phase charge state at pH 2 (E).**
